# Amino Acid Remodeling Precedes Malignant Transformation in Oral Epithelial Cells

**DOI:** 10.64898/2026.10.23.761404

**Authors:** Kaustav Das, Sharvari Haldavanekar, Deeptha Ishwar

## Abstract

**Background:** The accessibility of the oral cavity and the presence of clinically identifiable oral potentially malignant disorders provide an important opportunity for oral cancer prevention. However, current clinical and histopathological assessments cannot reliably predict which lesions will undergo malignant transformation. Here, we investigated amino acid alterations and related proteomic remodeling that characterize potentially malignant oral epithelial cells relative to healthy and malignant cells.

**Methods:** Healthy, potentially malignant, and malignant oral epithelial cell models were analyzed using Raman spectroscopy and liquid chromatography-mass spectrometry (LC-MS) based proteomics. Raman spectral profiles were compared across the three states of epithelial transformation, focussing on aromatic amino acid associated spectral regions. Proteomic changes were evaluated using differential abundance and pathway analyses, and the potentially malignant remodeling patterns were integrated with Raman findings.

**Results:** Raman spectroscopy identified distinct biochemical profiles across the three cellular states, with prominent alterations in the 1000–1165 cm⁻¹ region. Phenylalanine associated features near 1001–1004 cm⁻¹ were increased in the potentially malignant state and subsequently decreased in malignant cells. In contrast, tyrosine-associated band near 1164 cm⁻¹ were reduced in potentially malignant cells and increased in malignant cells. Proteomic analysis identified extensive stage associated remodeling of amino acid metabolism, transport, and protein synthesis. Early alterations included changes in GOT1, GOT2, GPT2, IVD, HIBCH, DBT, DLD, ALDH6A1, ALDH4A1, SHMT1, CTH, and other amino acid associated proteins. Integration of the datasets identified convergence between aromatic amino acid associated Raman features and early proteomic remodeling.

**Conclusions:** Potentially malignant oral epithelial cells exhibited distinct Raman and proteomic signatures that differentiated them from both healthy and malignant cells. The similarity between aromatic amino acid-associated Raman features and changes in proteins involved in amino acid metabolism and transport supports the presence of a distinct biochemical state associated with potential malignancy. These findings identify candidate molecular features for further validation in patient-derived lesions and future studies of malignant-transformation risk.

## 1 Introduction

Oral squamous cell carcinoma (OSCC) is the primary malignancy of the oral cavity and is associated with substantial morbidity and mortality, particularly when diagnosed at an advanced stage [1–5]. Surgical management results in extensive resection of oral and maxillofacial tissues, resulting in difficulties with mastication, swallowing, speech, facial appearance, and overall quality of life. Nevertheless, oral cancer presents an important opportunity for prevention and early intervention because the oral cavity is directly accessible to clinical examination. Visible mucosal alterations, frequently presenting as persistent white, red, or mixed red-and-white lesions, can be identified and monitored before invasive cancer develops [6–15]. These lesions are classified as oral potentially malignant disorders (OPMDs) and represent clinically detectable stages during which malignant transformation could potentially be prevented or identified earlier. OPMDs comprise a heterogeneous group of oral mucosal conditions associated with an increased risk of malignant transformation [16–20]. Although histopathological assessment of epithelial dysplasia continues to be the primary method of diagnosis, morphological grading does not consistently predict the progression of malignancy. Also, lesions with similar histological features often exhibits different biological behavior [21–26]. In fact, the term “oral potentially malignant disorders” was recommended in place of earlier terms such as “precancerous lesions” and “precancerous conditions” to recognize the uncertainty and heterogeneity of these disorders as only a subset undergoes malignant transformation, and the outcome of an individual lesion cannot be reliably predicted [27]. These limitations emphasize the need to identify molecular alterations that characterize the potentially malignant state before the onset of malignancy.

Metabolic reprogramming is a fundamental feature of cancer development, promoting proliferation, energy production and redox homeostasis [28–32]. Amino acid metabolism is a crucial process in the malignant transformation of epithelial cells because amino acids provide carbon and nitrogen for protein and nucleotide synthesis, contribute to tricarboxylic acid cycle, and participate in redox regulation and cellular signaling [33–40]. Metabolomic studies have identified amino acid-related alterations in saliva and other biological samples from patients with oral leukoplakia and OSCC, while proteomic studies have reported changes in proteins associated with glycolysis, oxidative stress, cytoskeletal organization, and other processes in potentially malignant oral lesions [36, 41–43]. Metabolomic and proteomic studies of OSCC have identified extensive alterations in amino acid-associated pathways with glutamate, methionine and other amino acids repeatedly implicated in disease [44–47]. Nevertheless, it remains unclear whether potentially malignant epithelial cells exhibit a unique biochemical phenotype that precedes malignant transformation or merely an intermediate form of the alterations observed in malignant cells. Furthermore, the relationship between changes in proteins involved in amino acid metabolism and transport and amino acid-associated Raman spectral features at the single-cell level remains poorly understood.

Raman spectroscopy provides a label-free approach for characterizing molecular composition through vibrational signatures, in a single cell level [48–52]. Raman spectra contain contributions from proteins, lipids, nucleic acids, carbohydrates and amino acid associated molecular groups [53–57]. Aromatic amino acids, particularly phenylalanine and tyrosine, contribute characteristic Raman features within the biological fingerprint region, including the prominent phenylalanine-associated band near 1001–1004 cm⁻¹ and tyrosine-associated band near 1164 cm⁻¹ [50, 57]. Raman spectroscopy therefore provides an opportunity to detect differences in the amino acid abundances accompanies epithelial alteration before they are morphologically evident [50, 54, 58–62]. Quantitative LC-MS based proteomics provides complementary information by measuring changes in protein abundance, including enzymes, transporters and other proteins involved in amino acid metabolism and nutrient uptake [63–66]. Comparing the findings from these two approaches can therefore help to determine whether amino acid-associated Raman alterations are accompanied by changes in proteins involved in amino acid metabolism and transport, thus strengthening the biological interpretation of our spectral findings **(Fig. 1)**.

**Fig. 1.**
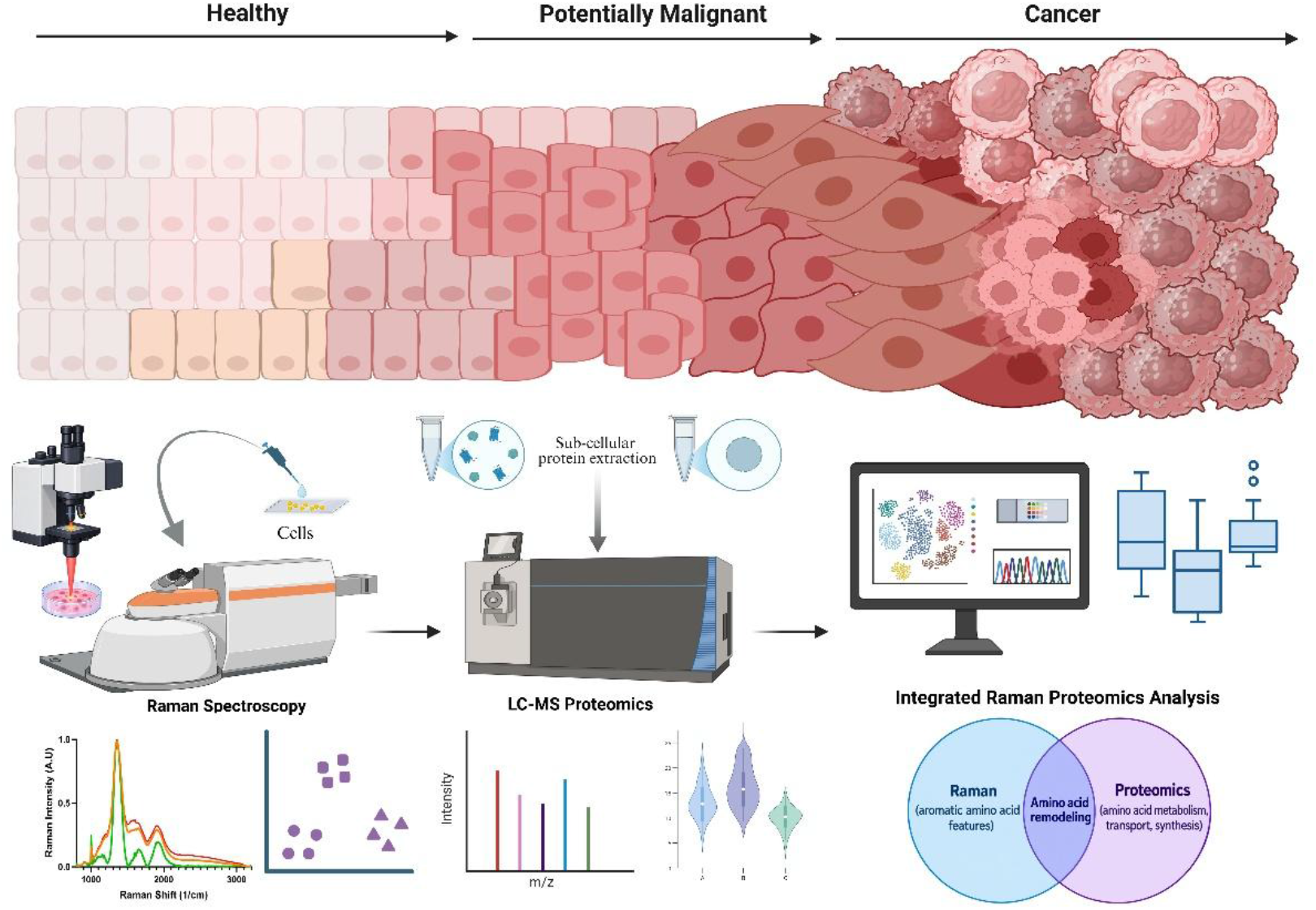
Integrated Raman-proteomic workflow for identifying early biochemical remodeling during epithelial transformation. The schematic illustrates the progression from normal oral epithelium through epithelial dysplasia, representing a potentially malignant state, to invasive oral squamous cell carcinoma. Healthy, potentially malignant, and cancer oral epithelial cells were profiled by Raman spectroscopy and quantitative LC–MS/MS proteomics. Raman analysis identified differential amino acid signatures, and proteomics characterized changes in amino acid metabolism, transport, and protein synthesis. Integration of both datasets identified amino acid remodeling as a distinctive feature of the potentially malignant state during epithelial transformation toward malignancy.

In this study, we first examined Raman spectral differences across the three cellular states and found that the potentially malignant cells showed particularly pronounced alterations within the 1000–1165 cm⁻¹ region, including changes near the phenylalanine-associated band at 1001–1004 cm⁻¹ and in regions with reported contributions from other aromatic amino acids. These alterations did not follow a simple progressive pattern from healthy to potentially malignant and then malignant cells. Rather, several spectral features were most prominent in the potentially malignant cells, suggesting that potentially malignant cells exhibit a distinct biochemical state. Quantitative proteomics identified 1,126 differentially abundant proteins among the 1,973 proteins quantified, including 1,029 proteins that changed between healthy and potentially malignant cells. Of these, 58 were increased and 971 were decreased in potentially malignant cells. Twenty-four differentially abundant proteins were associated with amino acid metabolism or transport, and 15 of these exhibited early remodeling in potentially malignant cells. This remodeling included increased GOT1 and reduced GOT2, GPT2, IVD, HIBCH, DBT, DLD, ALDH6A1, ALDH4A1, SHMT1, and CTH, indicating alterations in transamination and in branched-chain amino acid, proline, alanine, serine, glycine, cysteine, and methionine metabolism. Changes were also observed in the amino acid transporters SLC7A5 and SLC3A2 and in proteins involved in ribosomal function, translation, and aminoacyl-tRNA synthesis. Together, these findings show that potentially malignant cells undergo extensive biochemical and proteomic remodeling that is distinguishable from both healthy and malignant cells, rather than merely displaying an attenuated malignant profile.

## 2 Materials and methods

### 2.1 Cell culture

Human oral squamous cell carcinoma cells (UCSF-OT-1109) were obtained from the American Type Culture Collection (ATCC), Human oral leukoplakia cells (MSK-Leuk1) were obtained from Memorial Sloan Kettering Cancer Center and Human oral gingival epithelial cells (B11) kindly provided by the laboratory of Dr. Amel Ben Lagha, Department of stomatology, University of Montreal were used as cancer, potentially malignant and healthy epithelial cell model. UCSF-OT-1109 cells were maintained in DMEM supplemented with 10% fetal bovine serum (FBS) and 1% penicillin-streptomycin, as recommended by ATCC. MSK-Leuk1 cells were maintained in keratinocyte growth medium. Cells were maintained at 37 °C in a humidified atmosphere containing 5% CO₂. B11 human gingival cells were maintained in Dermal Cell Basal Medium supplemented with the ATCC Keratinocyte Growth Kit. Culture medium was replaced every 2–3 days, and cells were passaged at approximately 80% confluence. Cells were harvested during the logarithmic growth phase for downstream Raman spectroscopy and proteomic analyses.

### 2.2 Cytoplasmic protein extraction

Cytoplasmic protein fractions were isolated were isolated using the NE-PER™ Nuclear and Cytoplasmic Extraction Reagents (Thermo Fisher Scientific, Waltham, MA, USA; catalog no. 78833) according to the manufacturer’s protocol. The cultured cells were washed twice with cold phosphate-buffered saline (PBS) and lysed sequentially to release the cytoplasmic contents and subsequently solubilize the remaining nuclear pellet, yielding two distinct fractions per sample. The cytoplasmic fractions were aliquoted and stored at −80 °C until proteomic analysis.

### 2.3 Raman spectroscopy

Raman spectra were acquired from B11, MSK, and UCSF cells using a Renishaw inVia confocal Raman spectrophotometer equipped with a Leica DMI 6000 epifluorescence microscope. The spectrophotometer was calibrated using silicon chips with a characteristic Raman band at 520 cm⁻¹. Raman spectra were acquired from 10 cells of each cell type using a 785 nm laser over a spectral range of 400–3000 cm⁻¹, with an acquisition time of 10 s per spectrum. Raman spectra were preprocessed by baseline correction, normalization, and cosmic-ray removal.

### 2.4 Quantitative proteomic analysis

High-resolution liquid chromatography–tandem mass spectrometry (LC–MS/MS) was performed using an Orbitrap Fusion Lumos instrument operated in data-dependent acquisition (DDA) mode. The Proteomic analysis included three biological samples from each cellular states of healthy (n = 3), potentially malignant (n = 3), and cancer (n = 3). Sample loading, gradient elution, MS acquisition parameters, and instrument calibration were performed. Raw data files were processed using MaxQuant against the UniProt human reference database. Peptides and proteins were identified using a target-decoy strategy with a false discovery rate (FDR) of 1%. Label-free quantitative (LFQ) proteomic data were generated from cytoplasmic protein fractions obtained from healthy, potentially malignant, and cancer samples. Protein abundance data were imported into R (version 4.4.2) for downstream analysis.

### 2.5 Raman-proteomic integration

Raman spectroscopic and proteomic datasets were analyzed independently and subsequently integrated at the level of shared amino acid associated pathways and biological processes.

For Raman spectroscopy, one way analysis of variance (ANOVA) was performed for each Raman shift across healthy, potentially malignant, and cancer groups, followed by Benjamini-Hochberg (BH) correction for multiple testing. Raman features were classified as potentially malignant associated when FDR < 0.05 and the mean intensity in the potentially malignant group exceeded that of both healthy and cancer groups. Pairwise comparisons between potentially malignant and healthy groups and between potentially malignant and cancer groups were additionally performed using Wilcoxon tests with BH correction. Aromatic amino acid associated Raman features were identified by matching spectral shifts to predefined characteristic regions of phenylalanine, tyrosine, and tryptophan. An aromatic Raman score was calculated by extracting the corresponding Raman variables, applying z-score transformation, and calculating the row-wise mean across the matched features. Differences in the aromatic Raman score among groups were assessed using ANOVA and pairwise Wilcoxon tests with BH correction.

To characterize proteomic remodeling across the epithelial transition we evaluated differential protein abundance across three pairwise comparisons: healthy-to-potentially malignant (HP), potentially malignant-to-cancer (PC), and healthy-to-cancer (HC). Differential remodeling was identified by ANOVA with FDR < 0.05 and an absolute HP log₂ fold change (|log₂FC|) ≥ 1. To distinguish early alterations from changes associated with subsequent malignant progression, proteins were computationally classified according to their transition specific patterns as Early Potentially Malignant (FDR < 0.05, |HP log₂FC| ≥ 1, |PC log₂FC| < 1), Cancer Transition (|HP log₂FC| < 1, |PC log₂FC| ≥ 1), or Progressive Remodeling (|HP log₂FC| ≥ 1 and |PC log₂FC| ≥ 1). Proteins involved in amino acid metabolism and transport were subsequently identified based on functional annotation. Proteins showing significant remodeling during the HP transition were considered potentially malignant associated alterations.

Raman and proteomic datasets were then integrated at the biological process level by comparing potentially malignant associated Raman features with changes in proteins involved in amino acid metabolism and transport. Raman features and proteins were grouped according to their associated amino acid related functions, and patterns observed across the three cellular states were compared to determine whether spectral alterations were accompanied by corresponding changes in amino acid associated protein abundance. All the analyses were performed in R (version 4.4.2)

### 2.6 Statistical analysis

One-way ANOVA followed by Benjamini–Hochberg correction was used to assess differences in protein abundance and Raman features across healthy, potentially malignant, and cancer groups. Pairwise comparisons were performed using Wilcoxon tests with BH correction where indicated. Principal component analysis (PCA) and partial least-squares discriminant analysis (PLS-DA) were used to visualize separation among the three cellular groups. Hierarchical clustering and variable importance in projection (VIP) scores were used to identify features contributing to group separation. Figures were generated using GraphPad Prism and R with the ggplot2 package.

## 3 Results

### 3.1 Raman spectroscopy reveals distinct biochemical signatures of amino acid for potential malignancy

Biochemical remodeling accompanies epithelial transformation and may occur before the acquisition of an overtly malignant phenotype. Because Raman spectroscopy captures the collective molecular composition of individual cells, we used it to determine whether potentially malignant oral epithelial cells exhibit biochemical features distinct from those of non-malignant and malignant cells. Raman spectra were acquired from 30 individual cells with 10 cells from each of the three cell models. Each spectrum represents the average of three consecutive acquisitions. The strongest stage dependent changes occurred within the 1000–1165 cm⁻¹ fingerprint region. Variance analysis identified 10 Raman features that contributed substantially to spectral heterogeneity, centered at 1002, 1004, 1124, 1127, 1128, 1154, 1156, 1157, 1160, and 1164 cm⁻¹. Raman intensity at 1002 cm⁻¹ and 1004 cm⁻¹ corresponding to phenylalanine were increased in potentially malignant cells compared with healthy cells and subsequently decreased in cancer cells. Conversely, features centered at 1124–1129 cm⁻¹ corresponding to several C–H stretching and vibration associated protein features showed reduced intensity in potentially malignant cells, followed by increased intensity in cancer cells. The mean intensity at 1129 cm⁻¹ decreased in healthy cells and subsequently increased in cancer cells. A similar pattern was observed for the tyrosine associated feature at 1164 cm⁻¹, which showed a marked reduction in the potentially malignant state and increased intensity in cancer cells. The malignant cells exhibited an inverse spectral pattern, with a lower intensity of the phenylalanine-related peak at 1002–1004 cm⁻¹ and higher intensities at 1128 and 1164 cm⁻¹. Phenylalanine and tyrosine are essential components of protein synthesis and are interconnected through aromatic amino acid metabolism, while tyrosine additionally participates in signaling and other biosynthetic pathways. The broader C–H-associated features reflect changes in protein and lipid biochemical composition and therefore may represent alterations in cellular structural and metabolic organization. Multivariate analysis confirmed the separation of the three biological groups based on their Raman spectral profiles. PCA loadings revealed distinct positioning of the three groups along the principal axes of spectral variation, while PLS-DA further resolved the potentially malignant group from both healthy and cancer cells **(Fig. 2)**. The corresponding loading profiles indicate that specific Raman spectral regions contribute to this discrimination. Notably, the oral potentially malignant lesion displayed a spectral profile distinct from both healthy and cancer cells, indicating early biochemical remodeling rather than a linear transition toward malignancy. This distinct intermediate state suggests substantial metabolic and biosynthetic reorganization preceding malignant transformation.

**Fig. 2.**
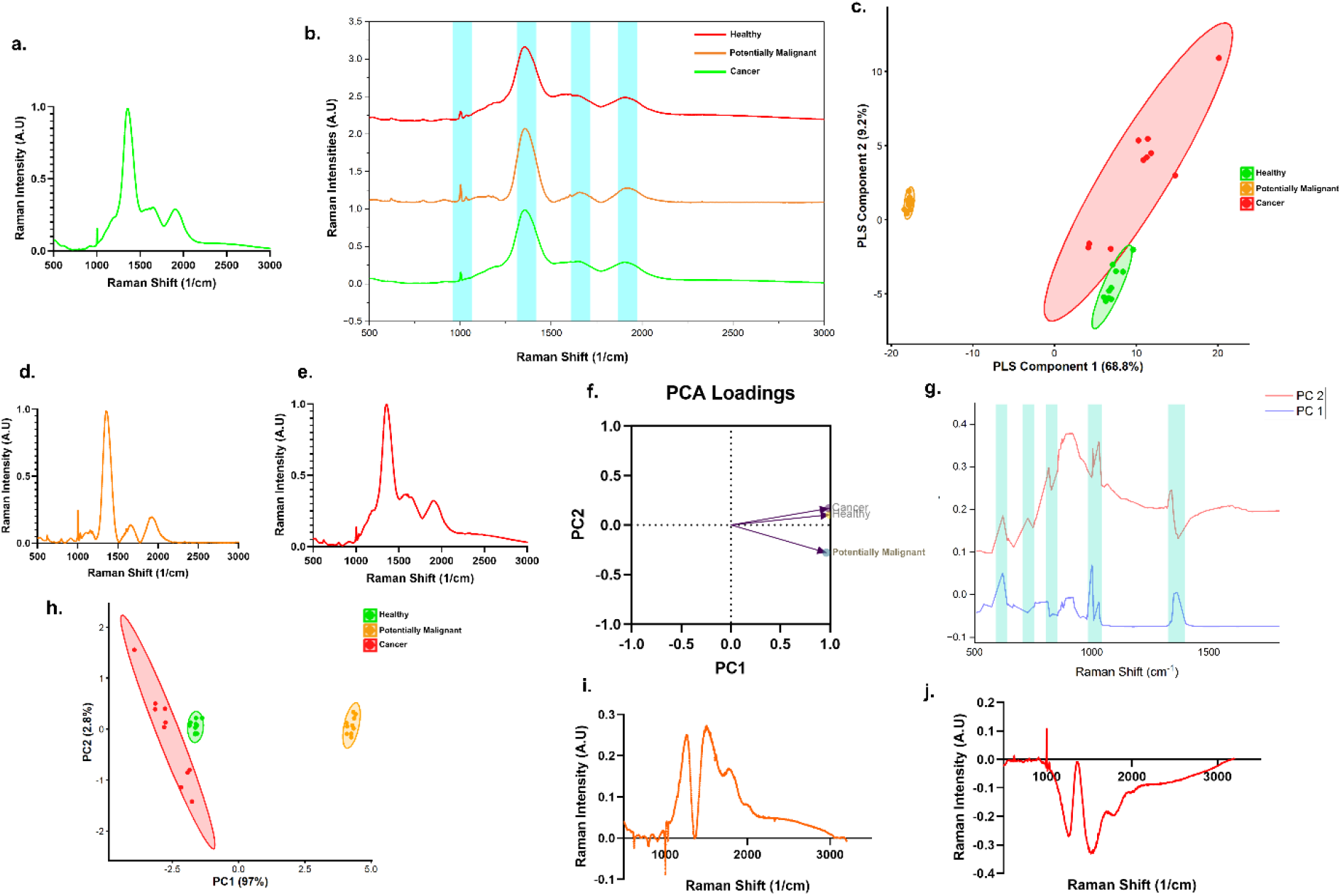
Raman spectroscopy reveals a distinct amino acid associated signature during epithelial transformation toward malignancy. **(a)** Mean Raman spectrum of healthy cells showing the overall cellular biochemical fingerprint across the measured spectral range (500–3000 cm⁻¹). **(b)** Baseline-corrected stacked Raman spectra across healthy, potentially malignant, and cancer cells highlighting differential spectral organization within the 1000–1165 cm⁻¹ fingerprint region. **(c)** Partial Least Squares Discriminant Analysis demonstrating separation of the three biological states based on global Raman spectral variation; ellipses indicate the 95% confidence regions for each group. **(d)** Mean Raman spectrum of potentially malignant cells showing a distinct biochemical spectral phenotype. **(e)** Mean Raman spectrum of cancer cells. **(f)** Principal Component Analysis (PCA) loading plot identifying Raman variables contributing to PC1 and PC2. **(g)** PCA loading profiles highlighting prominent spectral contributions near 1001–1004 and 1124–1164 cm⁻¹. **(h)** PCA score plot of top 10 Raman spectral features demonstrating separation of healthy, potentially malignant and cancer cells. **(i)** Differential Raman profiles highlighting the prominent 1001–1004 cm⁻¹ spectral feature associated with the potentially malignant state. **(j)** Differential Raman profiles highlighting the spectral features associated with cancer.

### 3.2 Proteomic profiling reveals early perturbations in amino acid associated metabolic protein abundances and translation factors

The Label free quantified proteomics data quantified 1,973 proteins after normalization and log2 transformation. Differential abundance analysis identified 1,126 proteins exhibiting significant variation across the three conditions (ANOVA, FDR < 0.05), indicating extensive remodeling of the cytoplasmic proteome during disease progression. The proteins involved in amino acid transport and metabolism showed pronounced changes during epithelial transition towards malignancy. CHORDC1, associated with protein folding, and CORO1C, involved in cytoskeletal organization and cell motility, were predominantly detected in cancer cells. Increased CHORDC1 reflects adaptation to the elevated proteotoxic stress generated by rapid proliferation, oncogenic signaling, and increased protein synthesis, thereby enhancing HSP90-dependent protein folding and stabilization of proteins required for cancer cell survival. Increased CORO1C promotes actin cytoskeletal remodeling, which helps in cellular motility and invasive behavior. In contrast, PSMD13 and CAND1, involved in protein turnover; FYCO1, associated with vesicular trafficking and autophagy; SUCLA2, ATP5I, and IDH3G, involved in mitochondrial metabolism; DGKA and ACSL3, associated with lipid metabolism and signaling; and GSTM2, GSTM3, and GSTM5, involved in redox regulation and detoxification, showed higher abundance in healthy cells and were reduced in potentially malignant cells.

Notably, GOT1 was strongly increased in potentially malignant cells compared to healthy cells, with further increase in the malignant state, which suggests that enhanced transamination and alanine, aspartate and glutamate metabolism are established early during transformation **(Fig. 3)**. In contrast, GOT2 was strongly reduced in potentially malignant cells compared with healthy cells and remained essentially unchanged between the potentially malignant and cancer states, which indicates an early alteration in TCA cycle. A coordinated reduction in IVD, HIBCH, DBT, and DLD in potentially malignant cells relative to healthy cells further indicated suppression of branched-chain amino acid catabolism, particularly valine, leucine, and isoleucine degradation, before the transition to the malignant phenotype. Similarly, ALDH6A1 and ALDH4A1 were reduced in potentially malignant cells compared with healthy cells, suggesting early remodeling of valine and proline metabolism. SHMT1 was also reduced in potentially malignant cells relative to healthy cells, indicating alteration of the serine and glycine. In addition, CTH and GPT2 were reduced in potentially malignant cells compared with healthy cells, implicating early changes in cysteine, methionine and alanine metabolism. These proteins showed little or no additional change between the potentially malignant and malignant states, indicating that their metabolic alterations were largely evident before malignancy. This pattern contrasts with proteins such as GLS, ASNS, ASS1, ALDH18A1, MTHFD2, and PYCR2, which underwent substantial changes between the potentially malignant and cancer states and therefore appear to be more closely associated with malignant progression. These findings indicate that substantial remodeling of amino acid metabolism occurs already in potentially malignant cells compared with healthy cells, characterized by increased GOT1 associated transamination besides reduced branched chain amino acid, proline, alanine, cysteine, and serine, glycine metabolic activities. These early metabolic alterations modify the availability and utilization of carbon and nitrogen substrates, which provides a potential metabolic flexibility and biosynthetic capacity that support subsequent malignant progression.

**Fig. 3.**
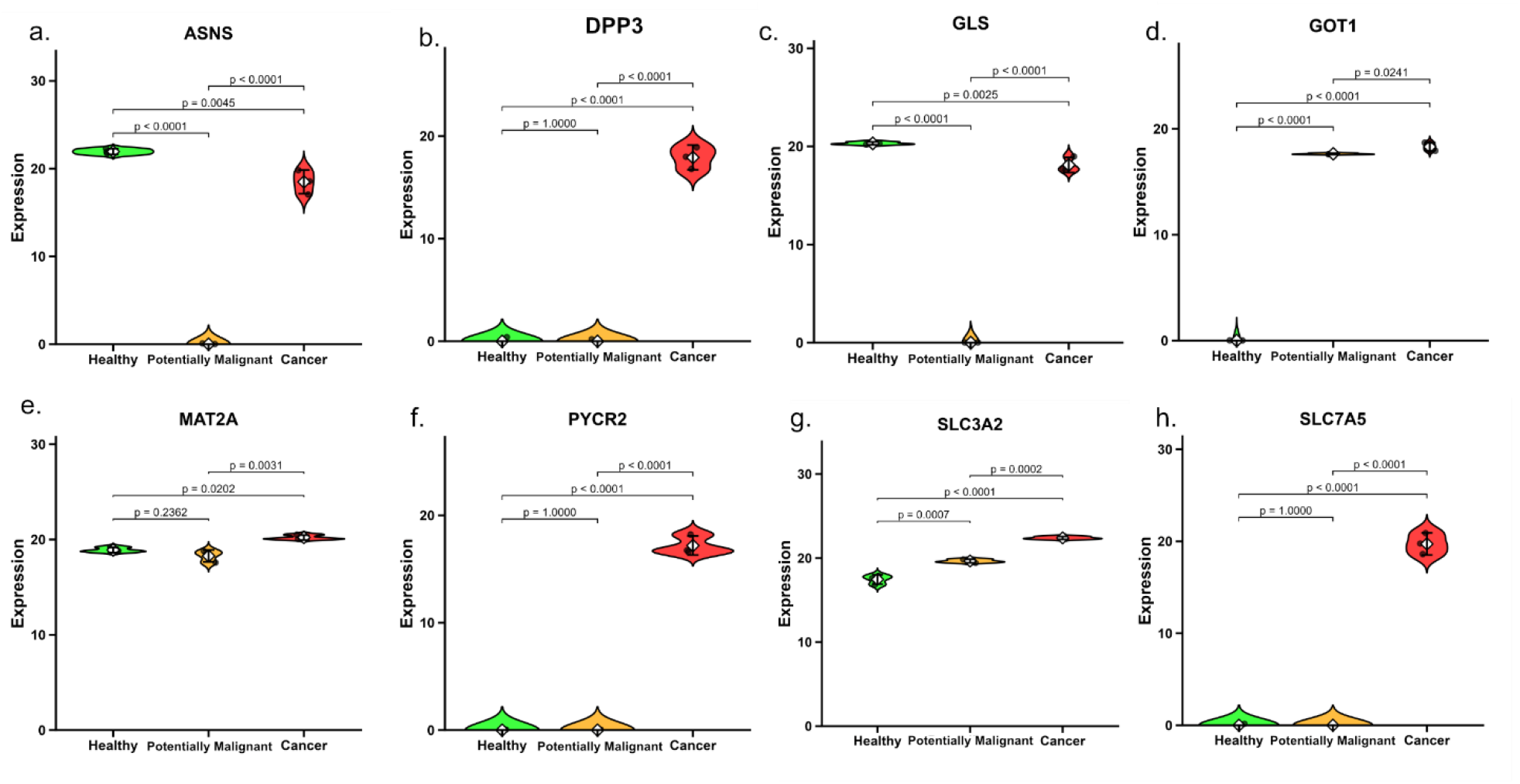
Proteomic profile reveals characteristic alteration of amino acid metabolism and transport during early epithelial transformation. **(a-h)** Abundance distributions of proteins involved in amino acid metabolism, biosynthesis, and transport during malignant transformation of epithelial cells. ASNS **(a)**, DPP3 **(b)**, GLS **(c)**, GOT1 **(d)**, MAT2A **(e)**, PYCR2 **(f)**, SLC3A2 **(g)**, and SLC7A5 **(h)** exhibit distinct abundance trajectories across healthy, potentially malignant, and cancer cells. Statistical comparisons are indicated above the distributions.

An extensive remodeling of the protein synthesis machinery was observed, with altered abundance of ribosomal proteins, translation initiation and elongation factors, and aminoacyl-tRNA synthetases. Multiple ribosomal proteins, including RPL5, RPL3, RPL8, RPL7, RPL13A, RPL35, RPL26, RPL17, RPL27, RPL6, and RPL10, together with translation factors such as EIF2S1, EIF2S2, EIF3B, EIF3F, EIF3H, EIF3I, and EIF4A1, exhibited malignancy associated changes in abundance. These findings indicate that alterations in amino acid metabolism in the potentially malignant model were accompanied by reorganization of the cellular protein synthesis machinery.

### 3.3 Integrated Raman-proteomics framework reveals coordinated amino acid remodeling in the potentially malignant state

Integration of the Raman and proteomic datasets revealed consistent patterns of biochemical remodeling across the three cellular states. Raman spectral analysis revealed an extensive biochemical remodeling across healthy, potentially malignant, and cancer cells. Amongst the 254 Raman variables analyzed, 218 had significant differences among the three cellular states (FDR < 0.05), and 75 features demonstrated their highest intensity specific to the potentially malignant state. To determine whether these spectral alterations were associated with aromatic amino acid associated biochemical signatures, predefined Raman regions corresponding to phenylalanine, tyrosine, and tryptophan were examined. From the 72 aromatic amino acid associated Raman variables, 18 exhibited their highest intensity in the potentially malignant state. A total of 12 features were associated with phenylalanine and 6 with tryptophan, whereas the tyrosine associated features were reduced in the potentially malignant state. The aromatic Raman score further demonstrated significant differences between potentially malignant and healthy cells (FDR = 0.0044) and between potentially malignant and cancer cells (FDR = 0.0211), supporting distinct biochemical remodeling of aromatic amino acids in the potentially malignant state **(Fig. 4)**.

**Fig. 4.**
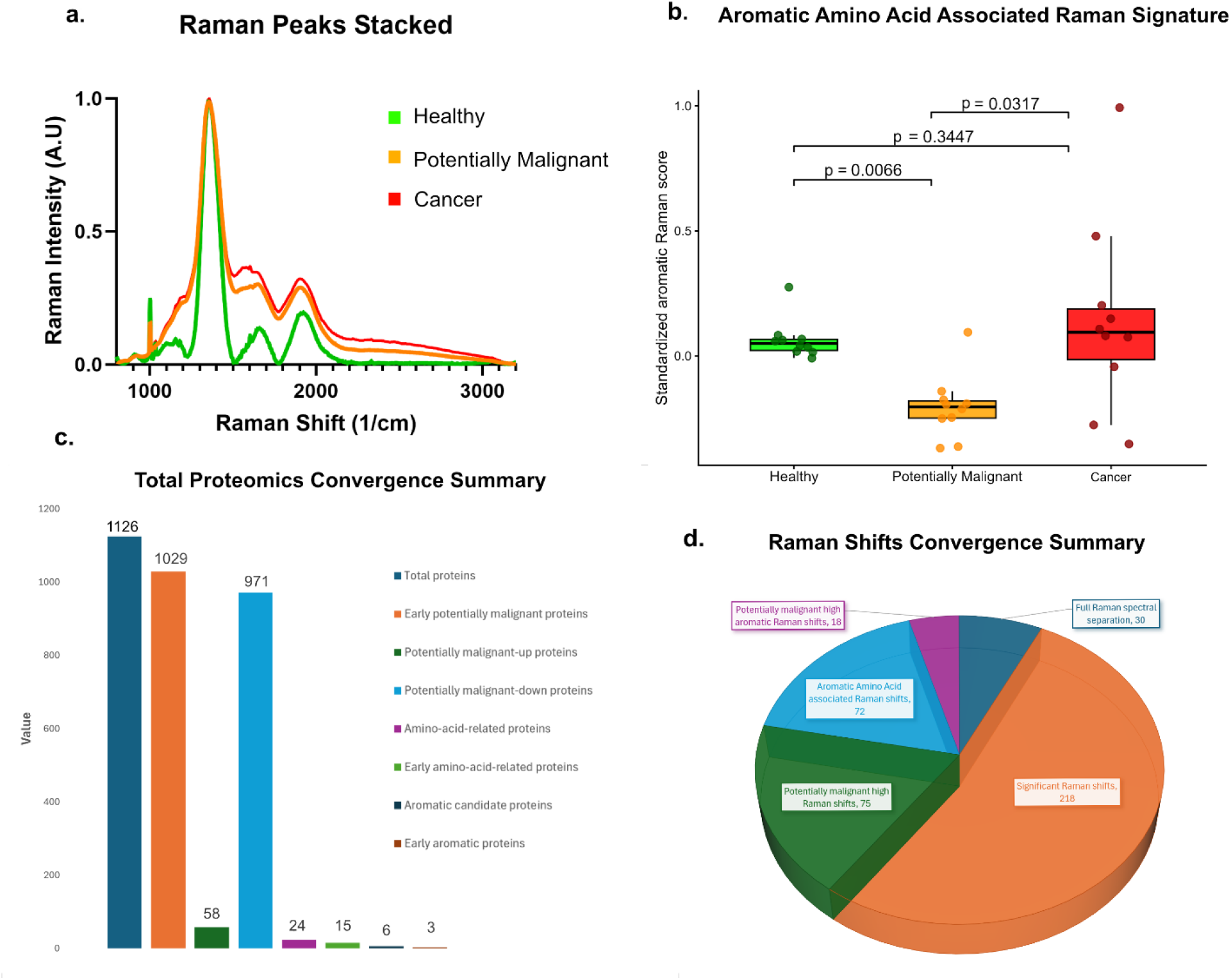
Integrated Raman-proteomics framework identifies early amino acid remodeling that differentiates the potentially malignant state from healthy and cancer cells. **(a)** Stacked mean Raman spectra of healthy, potentially malignant, and cancer cells showing distinct stage dependent spectral profiles, with prominent differences across the Raman fingerprint region. **(b)** Aromatic amino acid associated Raman signature showing the distribution of standardized Raman scores across healthy, potentially malignant, and cancer cells. The potentially malignant state exhibited a distinct aromatic Raman profile relative to both healthy and cancer cells. **(c)** Summary of proteomic remodeling across the three cellular states. Amongst 1,126 differentially abundant proteins, 1,029 exhibited significant healthy to potentially malignant remodeling, including 58 potentially malignant upregulated and 971 potentially malignant downregulated proteins; 24 amino acid associated proteins, 15 early amino acid associated proteins, 6 aromatic amino acid associated candidate proteins, and 3 early aromatic proteins were identified. **(d)** Of the 254 analysed spectral positions, 218 showed significant differences in intensity among the three cell states and 75 had their highest mean intensity in the potentially malignant cellular states. Among the 72 spectral positions located within predefined aromatic amino acid-associated regions, 18 had their highest mean intensity in potentially malignant cells.

Proteomic analysis showed a similar pattern of extensive remodeling during disease progression. Screening of amino acid associated proteins identified 24 proteins with significant changes, among which 15 exhibited early remodeling associated with potentially malignant cells. These changes included alterations in amino acid transport and metabolic proteins, which indicates that biochemical pathways involving amino acids are extensively remodeled during the early transition from healthy epithelium towards potential malignancy. Within the aromatic amino acid associated protein set, 6 proteins were identified, which are GOT2, GOT1, GPT2, SLC7A5, SLC3A2, and COMT. Three proteins GOT2, GOT1, and GPT2 showed early potentially malignant remodeling. GOT2 and GPT2 were reduced in the potentially malignant state, whereas GOT1 was increased and remained elevated in cancer cells. These findings suggest that oral potentially malignant lesions represent a distinct cellular state in which amino acid metabolism and biosynthetic processes are remodeled before malignancy develops.

## 4 Discussion

Oral epithelial transformation is a multistep process where non-malignant cells progressively undergo molecular and metabolic changes that may eventually facilitate invasive malignancy. The potentially malignant stage is particularly important because it represents a clinically detectable window during which progression may still be halted or detected before invasive cancer develops. However, this intermediate state is biologically heterogeneous with some lesions remaining stable whereas others progress despite having similar histopathological appearances. [67–69] Understanding the molecular features of potentially malignant epithelial cells is therefore essential for distinguishing potentially malignant state from both healthy epithelium and established cancer and for improving the assessment of malignant-transformation risk.

The principal finding of this study is that the potentially malignant oral epithelial cells exhibit a biochemical and proteomic profile distinguishable from both healthy epithelial cells and malignant cells. Importantly, the observed changes did not consistently follow a linear pattern from healthy/non-malignant to potentially malignant and then malignant cells. The 1001–1004 cm⁻¹ region exhibited the strongest Raman spectral variation. This is a characteristic region associated with the aromatic ring breathing vibration of phenylalanine. [50, 51] The raman peak intensity changes in these features therefore suggest remodeling of phenylalanine associated biochemical components during cancer progression. Additional alterations within the 1124–1164 cm⁻¹ region are consistent with changes in global protein component and aromatic amino acid associated vibrations, including contributions from tyrosine. The non-linear distribution of these spectral features supports the presence of a distinct biochemical organization in the potentially malignant cell model.

Proteomic analysis provided complementary molecular information. Of the 1,973 quantified proteins, 1,126 exhibited significant variation among the three biological states, demonstrating extensive stage dependent proteomic remodeling. Notably, proteins involved in amino acid transport and metabolism, including SLC7A5, SLC3A2, GOT1, DPP3, PYCR2, ASNS, MAT2A, and GLS, showed significant alterations across disease progression. Reduced abundance of IVD, HIBCH, DBT, and DLD suggested remodeling of proteins involved in branched-chain amino acid catabolism, while changes in ALDH4A1, ALDH6A1, SHMT1, and CTH implicated proline, valine, serine, glycine, cysteine, and methionine-associated processes. The marked increase in SLC7A5 and progressive elevation of SLC3A2 are particularly relevant to the Raman findings. SLC7A5, LAT1 forms a functional complex with SLC3A2, CD98 and transports large neutral amino acids, including phenylalanine and tyrosine. Increased LAT1 activity is widely associated with the enhanced amino acid requirements of cancer cells and metabolic reprogramming [41, 41, 44, 44, 70–74]. Thus, the proteomic remodeling of SLC7A5, SLC3A2 provides the molecular basis for the altered phenylalanine and tyrosine associated Raman signatures observed in the cancer cells.

The results show consistency between Raman derived biochemical phenotypes and proteomic alterations in amino acid metabolism and transport. Integration of Raman spectroscopy with proteomics provides the phenotypic information along with the molecular alterations, which strengthens its potential to identify biochemical changes associated with early malignant transformation. Phenylalanine, tyrosine and other amino acid associated Raman spectral alterations and changes in the amino acid transport proteins, provide evidence for studying potentially malignant state as a distinct condition for studying the progression of epithelial cells towards malignancy.

However, this study has some limitations. This study was conducted using in vitro cell line models to characterize biochemical and proteomic remodeling during oral epithelial transformation. Future studies should include patient-derived samples to validate the identified metabolic signatures. Additionally, targeted proteomics approaches should be employed to confirm and quantify specific amino acid metabolism and transport proteins identified in this discovery-phase study, thereby strengthening biomarker validation for potential clinical implementation.

## 5 Conclusion

In this study, we identified a distinct biochemical and proteomic signature associated with the potentially malignant state. Integrated Raman spectroscopy and quantitative proteomics prominent alterations in aromatic amino acid-associated spectral regions and changes in proteins involved in amino acid metabolism, transport, and protein synthesis. The distinct Raman spectroscopic and proteomic trajectories indicate that potentially malignant cells undergo a distinct metabolic and proteomic reorganization rather than representing a linear transition from healthy to cancer cells. These findings provide a basis for understanding the potentially malignant state as an early stage of epithelial transformation and support further evaluation of multimodal biochemical profiling for early detection of malignant progression.

## Data Availability

The data supporting the findings of this study are available from the corresponding author upon reasonable request.

## Code Availability

The code used to analyze the data and generate the results of this study is available from the corresponding author upon reasonable request.

## Clinical Trial Number

Not applicable.

## Author information

### Contributions

**Kaustav Das:** Conceptualization, methodology, formal analysis, investigation, writing – original draft. **Sharvari Haldavanekar:** Conceptualization, methodology, writing – review and editing. **Deeptha Ishwar:** Conceptualization, methodology, supervision, writing – review and editing and funding acquisition.

## Declaration

### Ethical Approval and Accordance

Ethical approval was not required, as the study did not involve human participants, human tissues, or animals. The work was conducted entirely using publicly available, de-identified database datasets and established in vitro cell line models and therefore falls under research that does not require formal ethics committee approval.

### Consent to participate

Not applicable.

### Consent to publish

Not applicable.

